# How fragile we are. Influence of STimulator of INterferon Genes, STING, variants on pathogen recognition and immune response efficiency

**DOI:** 10.1101/2021.07.12.452045

**Authors:** Jeremy Morere, Cécilia Hognon, Tom Miclot, Tao Jiang, Elise Dumont, Giampaolo Barone, Emmanuelle Bignon, Antonio Monari

## Abstract

The STimulator of INterferon Genes (STING) protein is a cornerstone of the human immune response. Its activation by cGAMP upon the presence of cytosolic DNA stimulates the production of type I interferons and inflammatory cytokines which are crucial for protecting cells from infections. STING signaling pathway can also influence both tumor-suppressive and tumor-promoting mechanisms, rendering it an appealing target for drug design. In the human population, several STING variants exist and exhibit dramatic differences in their activity, impacting the efficiency of the host defense against infections. Understanding the differential molecular mechanisms exhibited by these variants is of utmost importance notably towards personalized medicine treatments against diseases such as viral infections (COVID-19, Dengue…), cancers, or auto-inflammatory diseases. Owing to micro-seconds scale molecular modeling simulations and post-processing by contacts analysis and Machine Learning techniques, we reveal the dynamical behavior of four STING variants (wild type, G230A, R293Q, and G230A-R293Q) and we rationalize the variability of efficiency observed experimentally. Our results show that the decrease of STING activity is linked to a stiffening of key-structural features of the binding cavity, together with changes of the interaction patterns within the protein.

## Introduction

The defenses of evolved organisms, including humans, against pathogenic infection rely on finely-tuned biological machineries involving several cellular signaling mechanisms. The cyclic Guanosine mono phosphate-Adenosine mono phosphate Synthase–STimulator of IN-terferon Genes (cGAS-STING) pathway is a key player acting as a cytosolic DNA- or RNA-probe. After sensing the presence of exogenous genetic material it triggers the immune response through the production of type I interferon and cytokines.^1,2^ Indeed, the recognition of aberrant nucleic acid fragments in the cellular cytosol, such as those secreted by bacteria or resulting from viral infection, stimulates the cGAS enzyme, which produces cyclic guanosine adenosine monophosphate (cGAMP). Subsequently, cGAMP is sequestered by STING, inducing its activation and the final production of type I interferon and pro-inflammatory cytokines. These processes will also cause the promotion of downstream inflammatory signaling for the protection of uninfected cells and the stimulation of adaptive immune response.^3^ As a consequence, STING is known to play a crucial and sometimes contrasting role in different biological responses including antiviral defense,^3,4^ the mediation of tumor-suppressive and tumor-promoting mechanisms,^5–7^ autophagy,^8,9^ skin wound healing^10^ and auto-inflammatory diseases development.^11,12^ Its delayed activation might also be involved in severe COVID-19 outcomes.^13,14^ In this context, it has also been underlined that the over-stimulation of the cGAS-STING pathways leads to an inflammatory-like cytokine response, which is also strongly correlated with severe forms of the SARS-CoV-2 infection.^**?**^ Hence, the modulation of the cGAS-STING pathway and the related regulatory proteins provides suitable targets for the development of a wide variety of potential anticancer, anti-pathogen, as well as anti-inflammatory drugs and vaccines.^15–18^

As a matter of fact, the structure of the human STING protein has been entirely resolved and mechanistic hypothesis about its activation have been sketched so far.^19–21^ STING is a transmembrane protein, which is mainly localized in the endoplasmatic reticulum (ER) and is composed of two equivalent monomers. From a structural point of view one can distinguish a N-terminal transmembrane domain, having a high density of *α*−helices, a cytoplasm-exposed C-terminal domain, containing the cGAMP binding site, and a short linker region connecting the two domains - see Figure 1-A. Upon recognition and binding with cGAMP the C-terminal domain undergoes an important structural reorganization which ultimately results in the polymerization of different STING monomers, hence in the activation of the immune response. The cGAMP binding site is constituted by a pocket in the C-terminal domain which is surrounded by overhanging tails (lid regions), forming flexible random coils in the apo form that stiffen into *β*−sheets in presence of the ligand. Experimental and theoretical studies have stressed out the importance of Arg232, Arg238, Tyr167, Ser241, Thr263, Thr267 for stabilizing the ligand within the cavity.^20,22^ From a biochemical point of view STING polymerization, crucial for its full activation, takes place through the exposure of two cysteine residues in the linker domain which leads to the formation of a sulfur bond bridging the two monomers. The dimerization efficiency strongly depends on the solvent accessibility of these residues which are embedded in a rather flexible protein region, which may nonetheless assume an *α*−helix arrangement. Their solvent exposure is also modulated by the shielding effects caused by two disordered C-terminal tails whose conformation can be strongly affected by the ligand-induced structural transition further justifying the cGAMP-induced activation.

**Figure 1:**
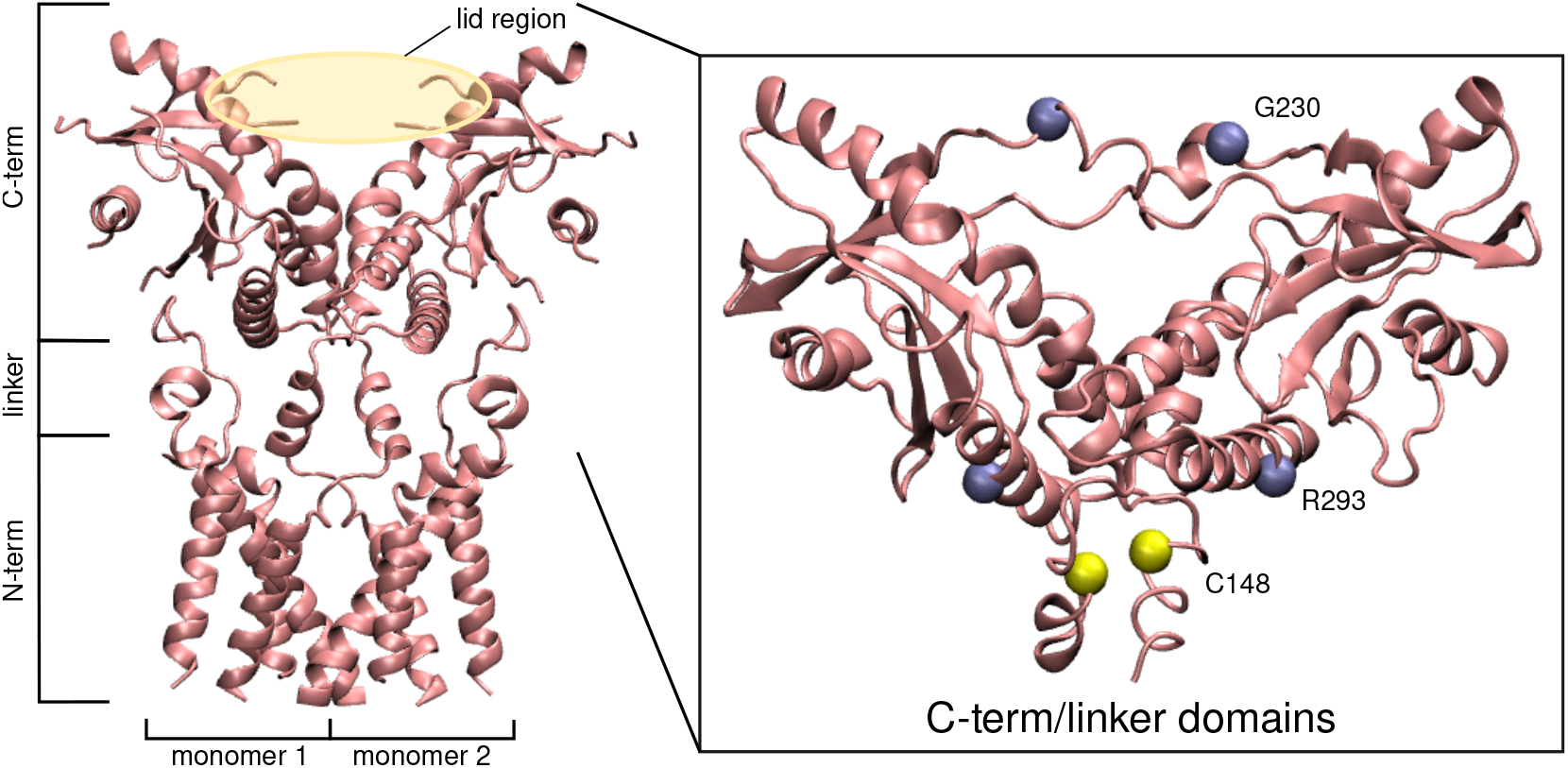
Cryo-EM structure of the full-length apo-STING dimer (PDB ID 6NT5). The magnified area shows the model used in out calculations, with the C-term and linker domains harboring reconstructed loops. The polymorphic sites and the cysteines involved in the multimerization are depicted as violet and yellow beads, respectively.

The STING-1 gene, also known as TMEM173, MPYS, MITA, ERIS, NET23, which is coding for the STING protein has a relatively high heterogeneity in the human population, which translates into the presence of several single-nucleotide polymorphisms involving the different domains of the protein. Albeit the R232 allele is the most common it only amounts to ∼60% of the general population and many variants co-exist: ∼20% of the population harbors the triple mutated R71H-G230A-R293Q (HAQ) variant, ∼14% carries the R232H polymorphism, ∼5% exhibits the G230A-R293Q (AQ) substitutions and ∼2% has a R293Q substitution.^3,23^ The activity of STING can differ from one variant to the other, with a possible loss-of-function associated with the HAQ and R232H variants and contrarily, gain-of-function with the N154S V155M V147L triple mutant, as found in patients suffering from vascular and pulmonary syndromes.^24^ Indeed, these combined differences induce variability in exogenous DNA- or RNA-sensing and consequently in the response to pathogen infections. Notably, the HAQ and R232H genotypes are associated with poor outcome in patients suffering from cervical cancer.^25^ Individuals carrying the HAQ polymorphism are more likely to contract the Legionnaires’ disease,^26^ probably more susceptible to infections, and less responsive to DNA vaccines.^27^ Interestingly, the loss-of-function held by the HAQ variant might be mostly attributed to the R71H and R293Q substitutions, while the G230A polymorphism would help maintaining partial response to bacterial cyclic dinucleotides.^28^ On the contrary, the R293Q substitution might provide enhanced protection against aging-associated diseases.^29^ However, gain-of-function variants might also contribute to auto-inflammatory diseases development.

On the bases of all these considerations, we used state-of-the-art all-atom molecular dynamics (MD) simulations to unravel at an atomic resolution the effects of common mutations (G230A, R293Q and G230A-R293Q see Figure 1) on the structural transition induced on STING by cGAMP binding. The biophysical knowledge so obtained was related to the observed differences in STING activation and hence immune response efficiency. In particular we focused on the effects of the mutations on either the exposition of the dimerization site or the accessibility of the binding pocket. To this aim we resorted to a truncated model of the protein involving only the cytoplasmatic C-terminal domain and the linker region. If this model, lacking the transmembrane domain, was not sufficient to fully describe the dimerization events, its reduced size allowed for a more extensive sampling of the rather complex interactions and long-range effects taking place upon cGAMP binding. Machine learning algorithms and contacts analysis were used to reveal both the key amino acids leading to the conformational transition and the allosteric consequences of the nucleotide mutations.

As a matter of fact, only few theoretical studies dealing with MD simulations of mutated STING have been reported so far. Notably, the study of the interaction of the DMXAA agonist on the mouse STING, and a cyclic dinucleotide (CDN) screening studies are available in the literature.^30**?** –35^ Our results provide for the first time a clear picture of the first steps of STING activation as well as its perturbation caused by common human variants such as the HAQ and AQ genotypes, which are associated to loss-of-function. More specifically, we pinpoint the role played by the global stiffening of the protein structure upon cGAMP recognition. Furthermore, we show how the combination of the different mutations involved in the HAQ variant leads to a drastic reorganization of the interaction network in the binding pocket that modulates the opening and closing of the protein, ultimately impacting cGAMP affinity and the immune response.

## Methods

### System setup

As mentioned in the introduction we have chosen a reduced model involving only the C-terminal domain of STING and the disordered linker region. Hence, we have excluded from our model all the N-terminal transmembrane domain. If this choice induces a drastic simplification of the model it also allows to take into account a reduced-size systems for which the statistical sampling will be deeper. It also allows to concentrate on the effects of the cGAMP binding and mutations, while neglecting the transmembrane effects. It is worthy mentioning though, that while our model is suited to explore the rigidification of STING and the differential effects of the mutants, it lacks an important element, i.e. the N-terminal disordered tails that protrudes outside the lipid membrane and are susceptible to interact with the linker region contributing to the modulation of the accessibility of the dimerization site. If we are aware of some of the biases induced by our choice it is important to underline that the disordered chains could be included only in presence of the full system, hence limiting the statistical sampling. Furthermore, their disordered nature will be particularly challenging to be captured with conventional force field that could lead to nonphysical over structuring of those domains. We should also stress out once more that in this contribution we mainly want to understand the effects of the STING mutations on the binding capability and on the linker domain structure.

For the apo systems, the starting structures were generated based on the C-terminal and linker domains from the cryoEM structure of the full-length human STING (PDB ID 6NT5^19^). The missing loops were reconstructed using SwissModel.^36^ The starting models for the systems with cGAMP were generated by homology to the cryoEM structure of the chicken STING harboring cGAMP (PDB ID 6NT7^19^), still with SwissModel. From these starting structures, the variants were built by mutating *in silico* the residues 230 and/or 293. Force field parameters for the cGAMP ligand were generated using the antechamber module of AMBER18^37^ for the derivation of RESP charges^38^ and the attribution of GAFF parameters^39^ - see parameters in SI. Standard STING residues were modeled using the ff14SB amber force field.^40^ The system was soaked in a cubic TIP3P water box with a 15Å buffer and potassium counter-ions were added to ensure a neutral total charge, resulting in systems of ∼135,000 atoms.

### Molecular Dynamics simulations

MD simulations were carried out using NAMD3^41^ for the dominant genotype of the human STING (wild type, WT) in its apo form and in the cGAMP-bound state. The Hydrogen Mass Repartitioning Method (HMR) was used to allow a 4 fs time step for the integration of the equations of motion. To prepare the system, 10,000 minimization steps were firstly performed imposing positional constraints on the protein backbone. Minimization run has been followed by 12 ns equilibration at 300K during which the constraints have been progressively released. The temperature has been kept constant using the Langevin thermostat with a 1.0 ps^*−*1^ collision frequency, electrostatic interactions were treated using the Particle Mesh Ewald (PME) protocol.^42^ After equilibration, the conformational ensemble was sampled along a 500 ns production run and structures were dumped every 40 ps. The same protocol was used for sampling three mutated states, involving the G230A (A-STING), the R293Q (Q-STING) and the G230A/R239Q mutation (AQ-STING), respectively. The starting protein structures were built manually by performing the point mutations from the WT system. Note that in the limit of our truncated system the AQ-STING can be considered of the highly spread and loss-of-function-inducing HAQ genotype.

### Structural Analysis

The cpptraj module of AMBER18^37^ was used to calculate distances, angles and root mean square deviations (RMSD) and to perform the clustering analysis. The latter was carried out according to deviations of the protein backbone and structures were clustered into 5 groups. The opening angle of the protein was computed as the angle involving the center of mass of the S162 residues lying at the bottom of the binding cavity, and the residues forming the *β*-sheet of the upper lobe of each STING monomer. The propensity of arginines to dive inwards the cavity was computed with respect to their distance to the S162 residues center of mass. Contacts analysis changes upon mutations of STING were computed using the GetContacts software (https://getcontacts.github.io/). Frequencies of contacts were calculated for each pair of residues and the most different patterns (75% threshold) identified among the STING variants were plotted as heatmaps using the ggplot2 package of R.^43^ Representations of the STING structure and projection of the contacts perturbation were rendered by VMD.^44^

### Thermodynamic integration

The perturbation of the free energy of binding upon mutation was assessed by Thermodynamic Integration. The soft core potential method was used to progressively alchemically mutate G230 to A or R293 to Q. The system being dimeric, each polymorphism implies two mutations in the system. To deal with it, we computed the delta G of binding by computing the thermodynamic cycle for one mutation on the first monomer, then a second thermodynamic cycle adding the mutation on the second monomer. Free energy calculations on the AQ double mutant were carried out from the A system, in two steps as well. 10,000 steps minimization, 60 ps-long thermalization and 1 ns production runs were performed with pmemd^37^ along 11 windows with lambda values varying from 0.0 to 1.0, and the convergence was further checked.

### Principle Components Analysis

Molecular dynamics simulation can provide important insight of the chemical and physical behavior of protein, however, the large dimensionality of the data obtained sometimes make it difficult to grasp the essence of the behavior of the model system. Principal components analysis (PCA) performs a linear mapping of the data to a lower-dimensional space by reconstructing a new configurational space that contains the most important degrees of freedom, providing a more intuitive way of understanding the chemical processes involved. In practice, PCA methods creates a covariance matrix from the coordinates of the trajectory, then computes its eigenvectors and corresponding eigenvalues. These eigenvectors serve as basis of a new configurational space, with each of them being a direction of motion. The first few eigenvectors with the highest eigenvalues are called Principle Components (PCs), and often contribute to the vast majority of the system’s behavior. In this paper, the sklearn library was used with a home brew script to perform PCA. We use the internal coordinates (inverse distance between geometric centers of two residues) of the trajectories as the input of PCA instead of the cartesian coordinates to provide better performance.^45^ The per residue importance were calculated by taking the sum of the weights of the PCs up to 80%, where the weight is defined as the eigenvalue of the corresponding PC over the sum of all the eigenvalues.

## Results and discussion

### STING structural features upon activation by cGAMP

#### Organization of the binding cavity

In the apo-state, STING experiences structural fluctuations between the open and close states, with the L225-N242 loops overhanging the binding cavity exhibiting a high flexibility - see Figures 1 and S1. Only D237 and R238 on the loops appear to be able to form stable interactions with K224, Y240, S241, N242, E260, or Y245 on the facing monomer. Yet, the loops remain disordered along the trajectory. The overall structure fluctuates between two opening states in the apo form, as evidenced by the two-maxima distribution of the opening angle shown in Figure 2 either around 110^*◦*^ or around 135^*◦*^. The angle is instead stabilized at 116.7 ± 0.1^*◦*^ by cGAMP binding leading to a stable close conformation. Upon binding of cGAMP, the two loops fold onto the ligand to structure into ‘lids’ at the edges of the binding cavity, as observed in previous experimental and theoretical studies.^19,20,22,46^ Two arginines belonging to the loops, (R238 on each monomer) dive towards cGAMP and get stabilized by very strong cation-*π* interactions with the purine moieties of cGAMP - see Figure 2 and Figure S2. The extremities of the loops forming the lid region get stabilized by interacting with the facing *α*−helix, which results in a very stable *β*−sheets conformation of the lid. At the extremity of both loops, D210 forms salt bridges with K236 and to a lesser extent K232, and hydrophobic interactions involving L180, A243, I245, P199 and V198 are also observed along the trajectory.

**Figure 2:**
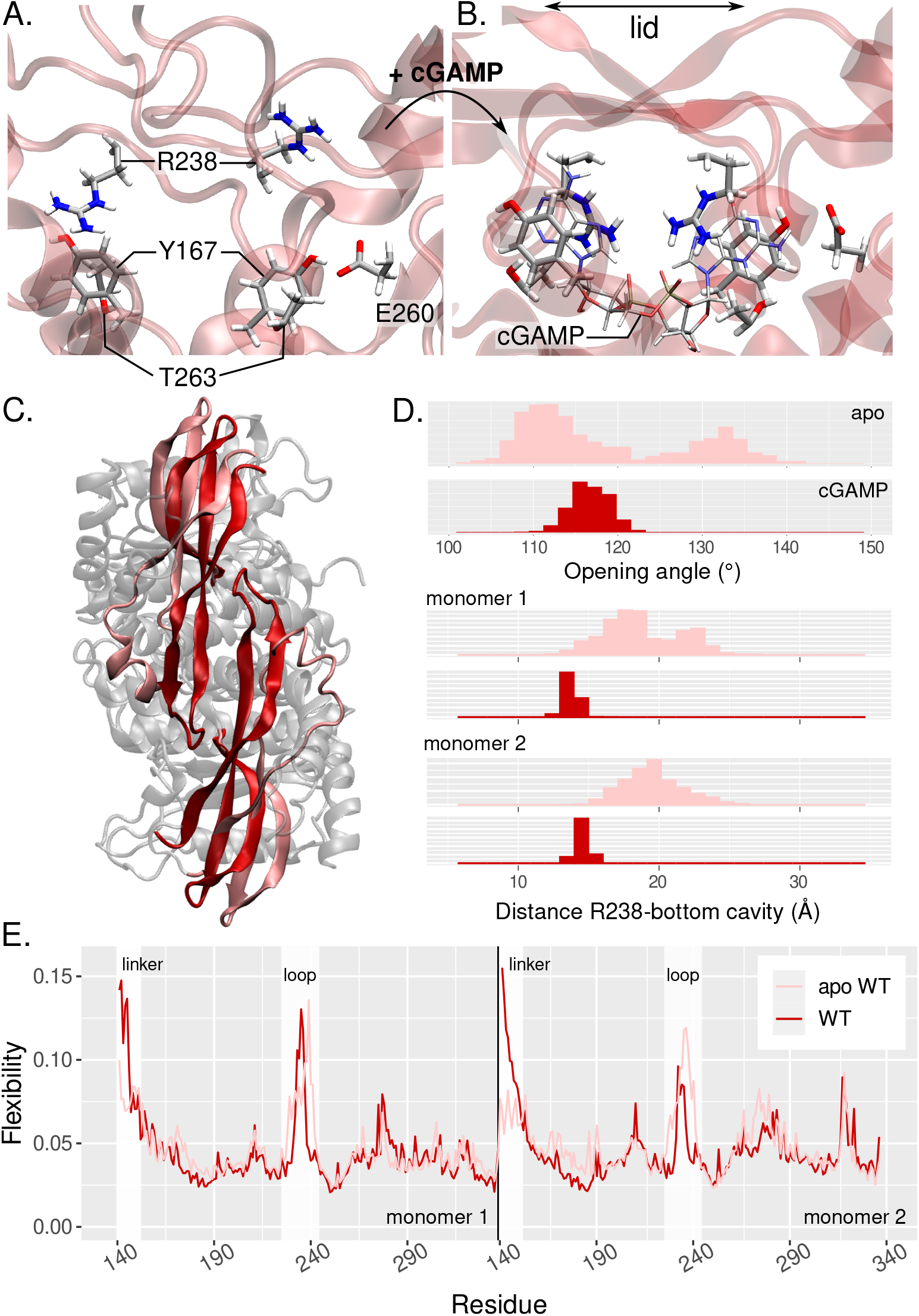
STING chemical and physical features perturbation upon cGAMP binding; light red coloration corresponds to apo-STING and dark red to STING bound to cGAMP. A. Representative structure of STING cavity without cGAMP and B. with cGAMP. R238, Y167, T263 and E260 of each monomer form the first coordination sphere around the ligand. B) Top view of the superimposed apo and complexed dimers. The lid region gets structured into beta sheets upon ligand binding. C) Distribution of the opening angle and of the distance between R238 and the bottom of the cavity. D) Flexibility profile of apo- and complexed-STING amino acids.

Globally the binding of cGAMP induces a very stable interaction network in the binding pocket. The latter involves, in addition to the R238 cation-*π* interactions already described, Y167 *π*-stacking with the cGAMP purines. Furthermore, hydrogen bonds between R238 and the ligand’s phosphates and between the nucleobases and E260 and T263 are also emerging - see Figure 2. These residues have been previously proposed to take part in cGAMP recognition^46^ and we also retrieve the previously-reported amino acids T267 in the second sphere of interaction together with Y163 and Y240 - see Figure S3. The R232 residues invoked in the literature^22^ is instead located on the external face of the lid and stays relatively far from cGAMP all along the trajectory in the WT, but, as it will be discussed in the following, it shows a more important role in the A and AQ variants. Interestingly, the S162 residue of both monomer, which lies at the bottom of the cavity and whose mutation to T or A destabilizes the cGAMP:STING complex,^46^ is also involved in the second sphere of interaction in our simulations.

#### Flexibility profile

In order to further probe the perturbation of the physical and structural properties resulting from the binding of cGAMP, we used a Machine Learning protocol based on principal component analysis (PCA) to post-process the MD trajectory and determine the flexibility profile of the protein. We successfully used this methodology on other DNA and proteic systems.^47–49^ The comparison of the WT STING flexibility profile in the apo and cGAMP-bound states underlines the stiffening of the lid region (residues 225-240) coupled to its structuring into a stable *β*−sheet as a result of the arginines diving towards the ligand - see Figure 2. Interestingly, one can instead distinguish an enhanced flexibility in the cytosol-transmebrane linker region, opposite to the binding pocket and harboring the cysteine residues that are involved in the disulfide bridge formation leading to the subsequent multimerization and to STING activation. The assessment of the cysteine exposure to the solvent also shows an increase of the number of water molecules around these residues in the bound state, suggesting that the residues are more accessible, hence more prone to encounter the reactive partners - see Figure S4. Nevertheless, as we used a truncated model in our simulations, this conclusion should be taken with some caution since we can not fully conclude that the same behavior would happen in the full-length structure. Yet the former stands as an interesting hypothesis on the allosteric regulation of STING activation that deserves to be investigated in further studies.

### Variant induced chemical and physical properties changes

#### G230A variant

Compared to the WT, the the open conformation of the apo state is favored by this variant as shown by the opening angle of 123.7± 0.4^*◦*^. Indeed, in this case only one single maximum in the distribution of the opening angle is observed. This fact is mainly due to a loss of contacts between the lid region and the facing monomer - see Figures 3 and S5. Upon binding of cGAMP, the structure closes around the ligand and the open angle drops to 115.1± 0.1^*◦*^, similarly to what is observed for the WT. The lid region gets highly structured and exhibits the lowest flexibility of all the variants, with an enhanced and contrasted interaction network with respect to the WT - see Figures 4 and S6. This observation is coherent with recent Isothermal Titration Calorimetry (ITC) results, which suggest that higher melting temperatures for the G230A cGAMP:STING complex might be related to a structural stabilization of the lid region, the ligand affinity of this variant being similar to the one observed for the WT.^46^ In line with these observations, the binding free energy change upon G230A mutation predicted by Thermodynamic Integration calculations is of only 1.20±0.25 kcal/mol - see Table 1.

**Figure 3:**
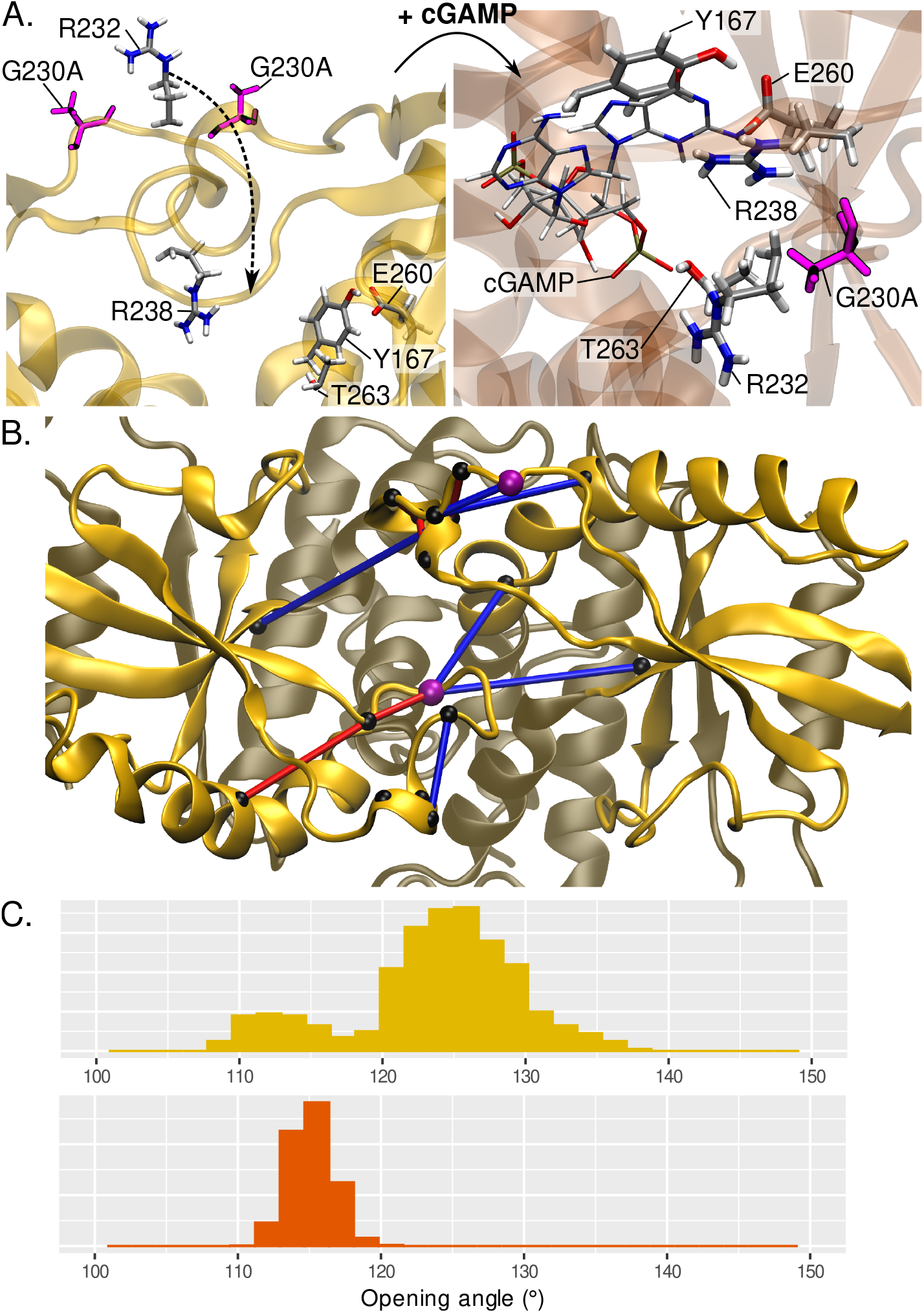
A. Structure of the A variant in the apo state. The mutated residues G230A of both monomers are depicted in purple, and the residues interacting with the guanosine moiety of cGAMP upon binding are displayed. On the right, zoom on the interactions with the guanosine within the binding cavity of the cGAMP-bound structure. R232 dives inwards to interact with the phosphate. B. Projection of the main changes of contacts in the lid region of the apo STING upon G230A mutation. Loss and gain of contacts are represented as blue and red tubes, respectively. The amino acids involved in the perturbed contacts are depicted as black beads and the G230A mutations as purple beads. C. Opening angle distribution of the apo and cGAMP bound states of A-STING.

**Figure 4:**
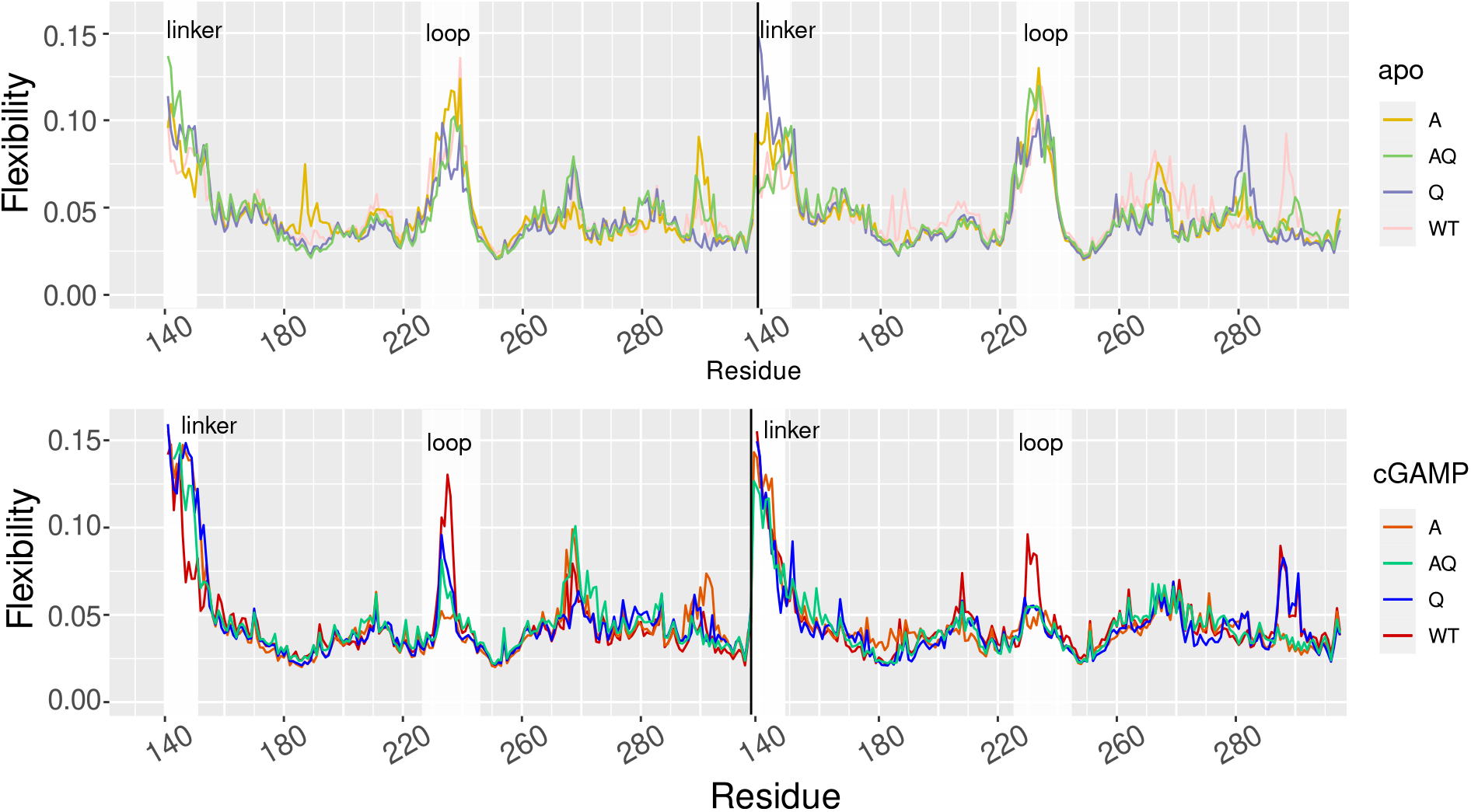
Flexibility profile of the four variants in the apo (top) and cGAMP (bottom) states.

**Table 1:** Relative free energy of binding (ΔΔ*G*) in kcal/mol upon mutation of the WT STING into A, Q, and AQ models as computed by Thermodynamic Integration calculations.

| Variant | $\Delta\Delta G$ | error |
| --- | --- | --- |
| G230A | 1.20 | $\pm 0.25$ |
| R293Q | 1.11 | $\pm 0.65$ |
| G230A-R293Q | 9.40 | $\pm 0.68$ |

On the contrary the strongly enhanced structuration of the lid region upon cGAMP binding is well evidenced by the flexibility profile - see Figure 4. The lid stiffening impacts the binding site organization around cGAMP. On the guanosine side of the ligand, the G230A mutation hampers the interaction of R238 with cGAMP. R238 is pushed further from the purine than in the WT, yet it is proximal enough to interact with the phosphate by hydrogen bonds. Interestingly, the nucleobase position is mainly maintained by *π*-stacking with Y167 and hydrogen bond with E260. Contrary to the WT, R232 dives towards the ligand to persistently interact with the phosphate group. T263 also interacts with the latter instead of the nucleobase in the WT structure. On the adenosine side of cGAMP, one retrieves the nucleobase *π*-stacking with Y167 and hydrogen bond with T263, as well as the interaction between R238 and the phosphate, although R232 is again further from the nucleobase than what is found for the WT-STING, preventing cation-*π* interactions - see Figure 3. Altogether, the influence of G230A mutation on STING function can be associated with the favoring of an open conformation in the apo state which is ideal to assure recognition and binding of cGAMP. Although, the interaction network in the binding pocket is altered, the ligand is still stabilized and the same increased flexibility and cysteine exposure in the linker region is observed - see Figure S4. Hence, one can conclude that the presence of the G230A mutation should favor global STING activation.

#### R293Q variant

Differently from the previous case, the R239Q mutation is located further from the binding pocket. Here, the Q-STING binding site harbors cGAMP in a similar fashion to the WT - see Figure S7. Both R238 residues interact through cation-*π* interactions and hydrogen bonds with the nucleobases and the phosphates of cGAMP. E260, Y167 and T263 also participate to the interactions network within the cavity, and the R232 residues remain in the second sphere of interaction yet pointing towards the bulk - see Figure S3. Interestingly, in the apo state, both R238 form very stable hydrogen bonds with the E260 on the facing *α*helices, which promotes a closed conformation characterized by an opening angle of 116.0± 0.2 ^*◦*^. The more moderate opening of the binding pocket might disfavor cGAMP access and hence its recognition - see Figure 5. Besides, the contact map underlines, for Q-STING, much more frequent interactions within the lid region itself but also with the surroundings spanning the residues 225 to 240 region in both monomers. Interestingly, these contacts are more pronounced than for the other variants and exhibit a contrasted pattern compared to WT-STING - see Figure S5. Conversely, cGAMP binding still induces a considerable enhancement of the flexibility of the linker regions, hence a more solvent-exposed cysteine. As a consequence, and in particular due to the promotion of a more closed conformation in the apo state that is susceptible to strongly perturb the recognition of cGAMP, this mutation should be correlated to a loss of activity of STING.

**Figure 5:**
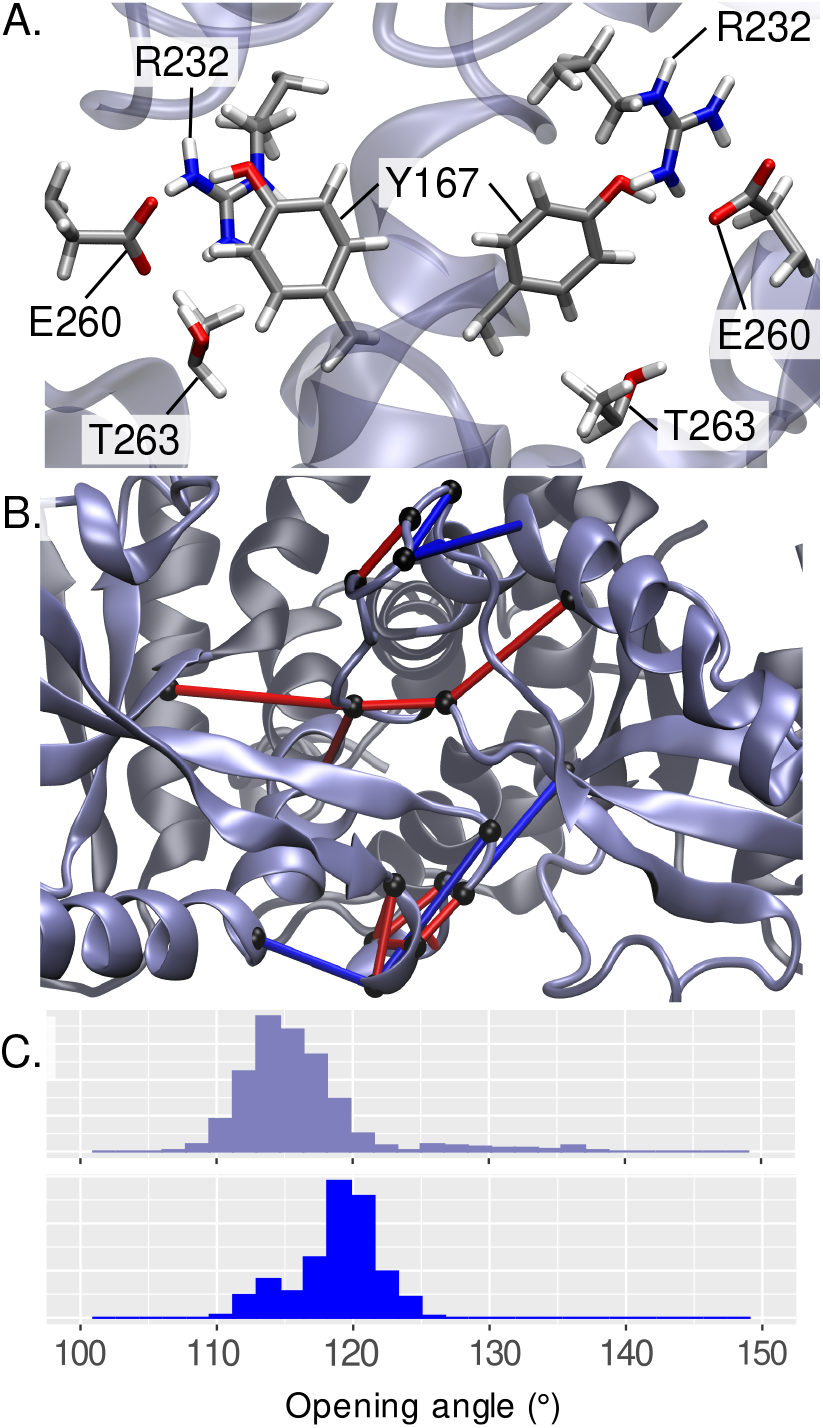
A. Organization of the binding cavity in the apo variant R293Q. B. Projection of the main changes of contacts in the lid region of the apo STING upon R293Q mutation. Loss and gain of contacts are represented as blue and red tubes, respectively. The amino acids involved in the contacts are depicted as black beads. C. Opening angle distribution in the apo (top) and cGAMP-bound (bottom) states of Q-STING.

#### G230A-R293Q variant

The AQ model, which may be directly related to the loss of function HAQ STING genotype, presents an organization of its binding site much alike the A variant. In the apo form, Y167 is closed to E260 an T263, while R238 remains in the cavity and R232 is in the bulk.

Upon ligand binding, R232 dives towards the cavity and interacts with the phosphate - see Figure 6. Contrary to the A variant, the *π*-stacking interaction with the guanosine is not disrupted, and R238 does not enters the cavity but rather interacts on the side. We retrieve here the interaction between cGAMP and E260, Y167 and T263. Importantly however, the opening angle distribution is centered around 122.4±0.1^*◦*^ and 113.9±0.1^*◦*^ in the apo and bound states, respectively. Interestingly, the opening angle for the AQ variant lies in between the open and closed conformations observed in the other variants, and it exhibits the lowest value upon ligand binding. Like the A and Q variants, a drastic stiffening of the lid region is still observed in passing from the apo to the bound states (see Figures 4 and S8), correlated with changes of contacts in this region of the protein which induce a higher structuration of the *β*−sheet lid - see Figures 6, S5 and S6. Interestingly, the relative energy of binding predicted by Thermodynamic Integration calculations indicates a strong perturbation leading to a lowering of the binding free energy of 9.40 ± 0.68 kcal/mol upon the AQ double mutation. Therefore, the loss of activity of the AQ (or HAQ) STING might arise from both a lower affinity as well as a lower accessibility to the binding cavity due to the stiffening of the protein structure and the more closed apo conformation.

**Figure 6:**
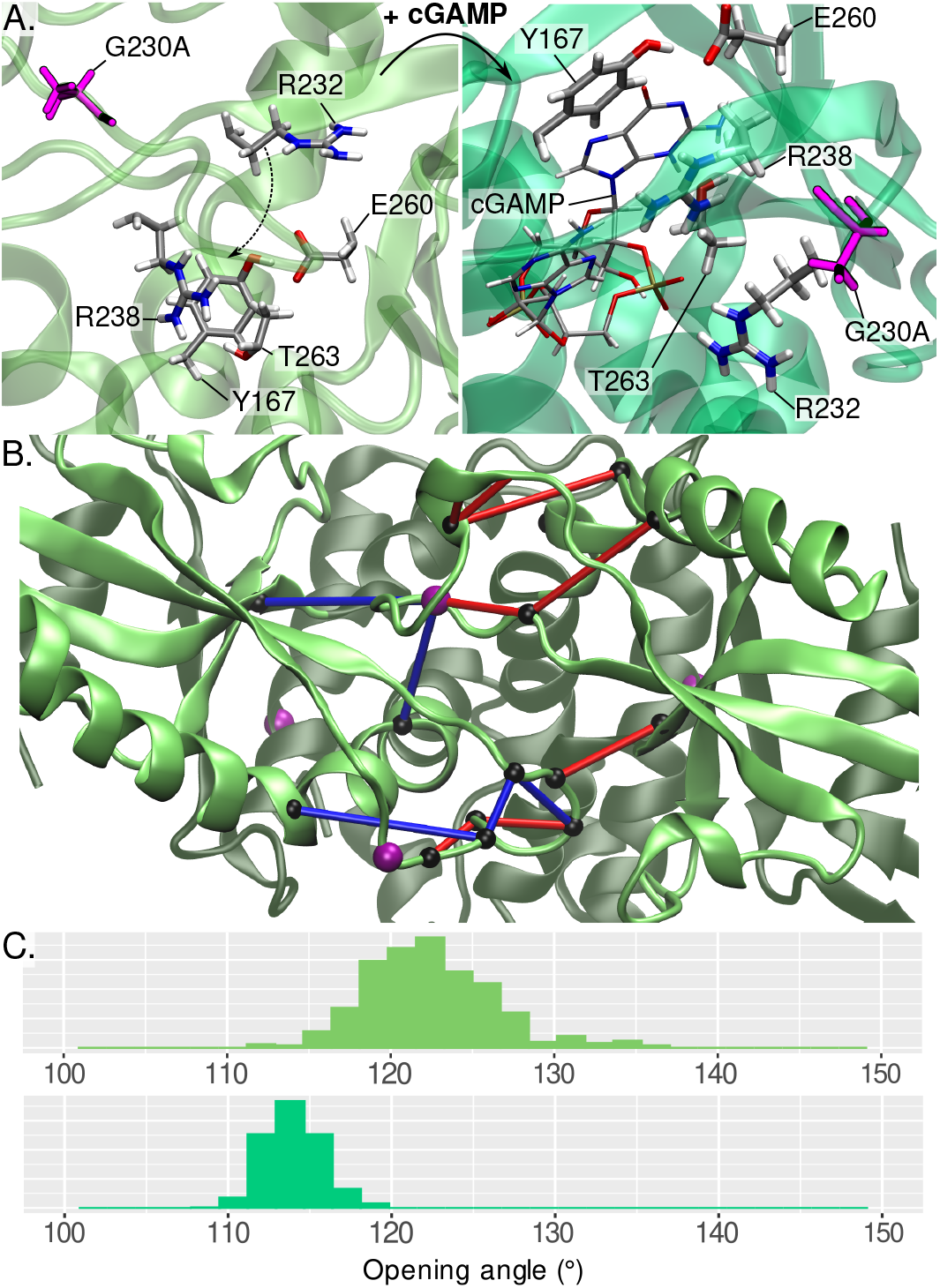
A. Organization of the binding cavity in the apo G230A-R293Q variant. The mutated residues G230A are depicted in purple, and the residues interacting with the guanosine moiety of cGAMP upon binding are displayed. On the right, zoom on the interactions with the guanosine within the binding cavity of the cGAMP-bound structure. B. Projection of the main changes of contacts in the lid region of the apo STING upon the G230A-R293Q double mutation. Loss and gain of contacts are represented as blue and red tubes, respectively. The amino acids involved in the contacts are depicted as black beads. C. Opening angle distribution in the apo (top) and cGAMP-bound (bottom) states of the AQ-STING.

## Conclusions

STING is a crucial transmembrane protein present in the cellular ER and involved in sensing exogenous genetic material and in triggering the immune response through proinflammatory pathways. Recently STING has also been associated to the cytokine storm which may lead to severe COVID-19 cases upon SARS-CoV-2 activation. In this contribution, by using high-level all atom MD simulations coupled with machine learning analysis, we have contributed to shed light on the fundamental mechanisms of STING activation. More particularly STING coding gene exhibiting various polymorphisms, we have also analyzed the effects of common variants on its structural transitions and ligand binding capability, with the aim to rationalize the loss of activity of some common variants. We have clearly seen that the interaction with cGAMP leads to an important remodeling of the interaction network of the protein, whose more important effect is the stucturation of the disordered loops overhanging the binding pocket into a lid region assuming a *β*−sheet arrangement. In the WT apo form the loops coexist in a closed and open conformation, characterized by different opening angles. Obviously an open conformation is necessary to allow the entrance of cGAMP into the binding pocket and hence its recognition. Interestingly, as revealed by our PCA-based machine learning analysis, the structuration upon cGAMP binding is also accompanied by a noticeable increase of the flexibility of the linker domain involved in STING activation via multimerization through the formation of sulfur bridges. Indeed the cysteine residues present in the region are much more solvent-exposed as an effect to cGAMP binding, hence favoring the probability of encounter other cysteines, hence of reaction. This long-range modulation is most probably at the base of the activation mechanism of STING, even if care should be taken to avoid over interpretation of our results due to the use of a truncated model missing the transmembrane domain.

As concern the role of mutations, contrasting effects have been evidenced depending on the specific mutation. However, we may recognize a remodeling of the protein rigidity profile and internal long range communication pathways. Indeed, changes in the lid region flexibility induce a perturbation of the interaction pattern within the cavity. Contrary to the A230G mutation, the R293Q mutation induces a stiffening of the lid region in both Q and AQ models, which in turn translates in a prevalence of the closed conformation in the apo form. As of note, while the tightening of the access to the binding pocket is strongly reduced in A-STING, the simultaneous presence of the two point mutations in AQ-STING leads to an intermediate situation compared to the WT. This in turn can be related to the observed loss of efficiency of AQ-STING, which in its HAQ form is present in about 20% of the global population. Indeed, the more difficult access to the binding pocket may lead to a less efficient recognition of cGAMP, and hence lower STING activation. Noteworthy, and despite strong remodeling of the interaction network in the lid region, the estimated binding free energy of cGAMP is only negligibly affected by the G230A mutation, while a significant increase is observed in case of the AQ model.

Our study allows, for the first time, a rationalization of the role of variants in STING contrasted phenotype, and more specifically highlights the role of long-range communication and the modulation of the prevalence of the open and close conformation in cGAMP recognition and STING activation. In the future we plan to extend the study in two directions: from the one side we will focus on other variants and isoforms, present in the general population and leading to constitutive over activation of STING, which may be of interest in the treatment of autoimmune diseases.^24,50^ On the other side, we will also complexify our model to introduce the transmembrane domain and a lipid bilayer to take into account all the possible alterations of the communication network and better rationalize the enhancement of the flexibility of the linker region upon cGAMP binding. Resorting to coarse grain approaches will also allow the explicit study of STING multimerization, as well as the effect of post-translational modifications, such as phosphorylation or ubiquitination, which may modulate STING activity.^51–53^ A most interesting future research line would also be the study of the interaction with viral proteases, present for instance in Zika or Dengue viruses, which reduce the host immune response by leading to the cleavage of STING.^54,55^ Ultimately, the enhanced comprehension of the basic mechanisms of STING activation may lead to significant advancements in the modulation of immunotherapeutic strategies, or in the development of host-targeted antiviral treatments.

## Supporting information

cGAMP force field parameters

## Acknowledgement

The authors are grateful to the GENCI HPC for computational resources on the IDRIS Jean Zay cluster under the project “Seek&Destroy”. Part of the calculations have been realized on the local LPCT computing ressources and on the regional Explor computing center in the framework of the project “Dancing under the light”. C.H. and T.M. thank the French and the Italian Ministries of Research, respectively, for their PhD fellowships. E.B. thanks the CNRS and French Ministry of Higher Education Research and Innovation (MESRI) for her postdoc fellowship under the “GAVO” program.

## Supporting Information Available

- Supplementary figures and tables
- cGAMP parameter files (CMP.zip)

