## Supplementary Materials for "How fragile we are. Influence of STimulator of INterferon Genes, STING, variants on pathogen recognition and immune response efficiency"

‡*Department of Biological, Chemical and Pharmaceutical Sciences. Universita degli Studi di Palermo, via delle Scienze 90126 Palermo, Italy.*

¶*Université de Lyon, ENS de Lyon, CNRS UMR 5182, Laboratoire de Chimie, F69342, Lyon, France*

§*Institut Universitaire de France, 5 rue Descartes, 75005 Paris, France*

∥*Université de Paris, CNRS, ITODYS, F-75006, Paris, France*

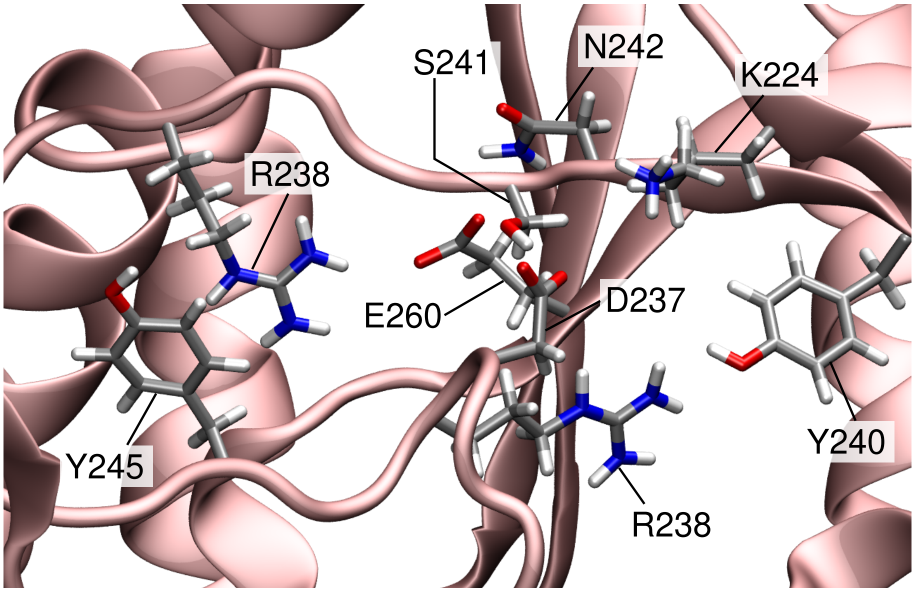


Figure S1 – Amino acids forming interactions in the lid region of the WT apo-STING.

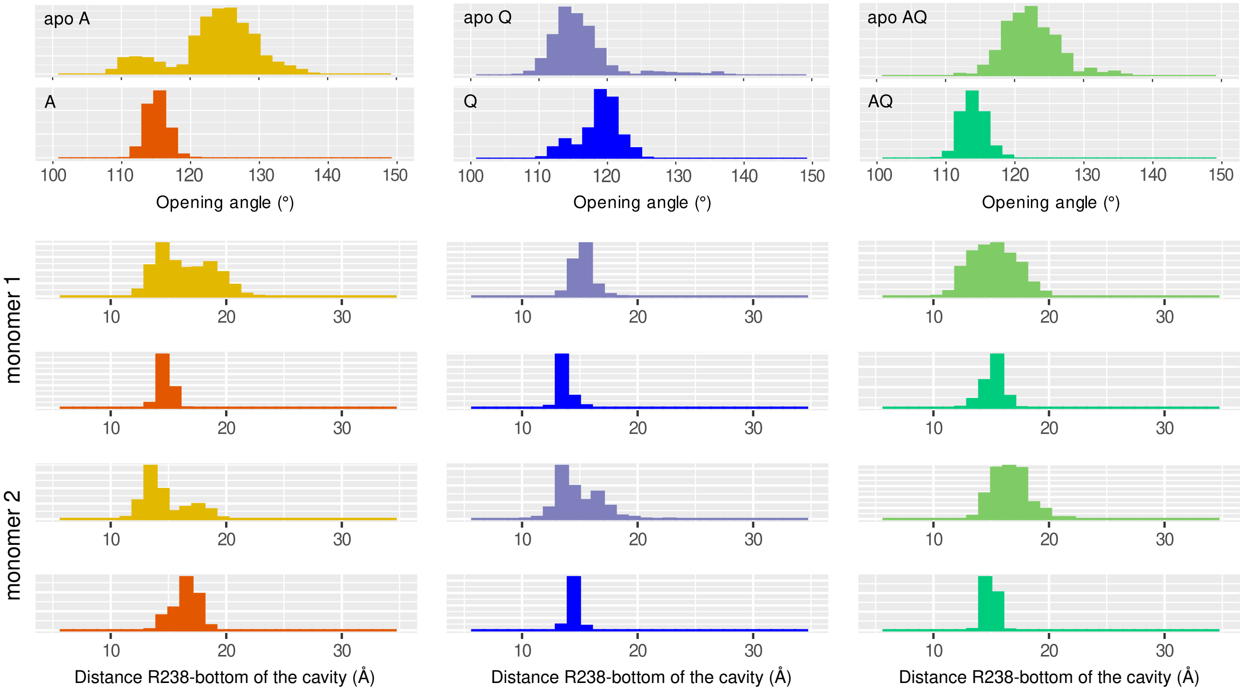


Figure S2 – Opening angle and distance R238-bottom of the cavity distribution for the A (left), Q (center), and AQ (right) variants in the apo (yellow, iceblue, lime) and the cGAMP-bound (orange, blue, green) states.

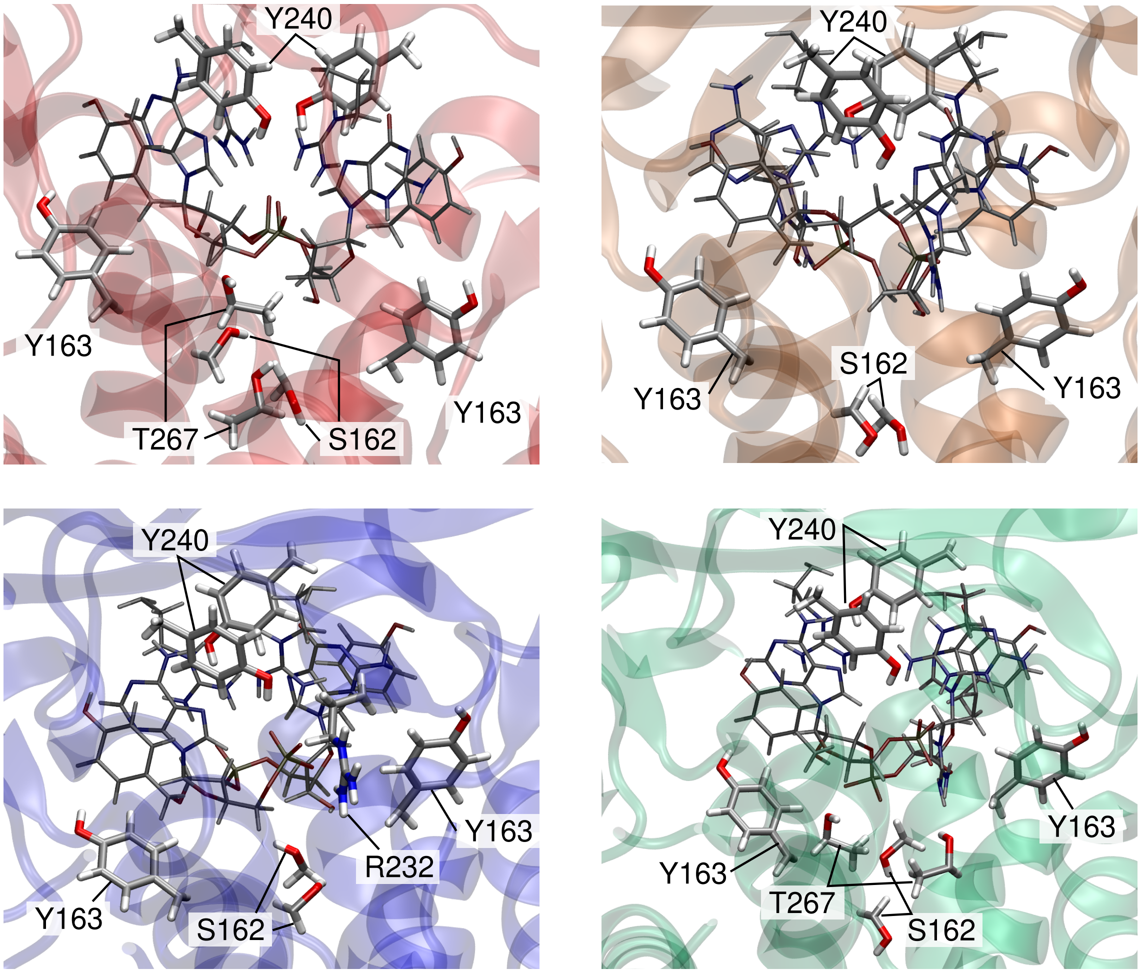


Figure S3 – Amino acids in the second sphere of interaction around cGAMP in each variant: WT (top left), A (top right), Q (bottom left) and AQ (bottom right). The ligand and R232 and Y167 are depicted in thiner and darker lines.

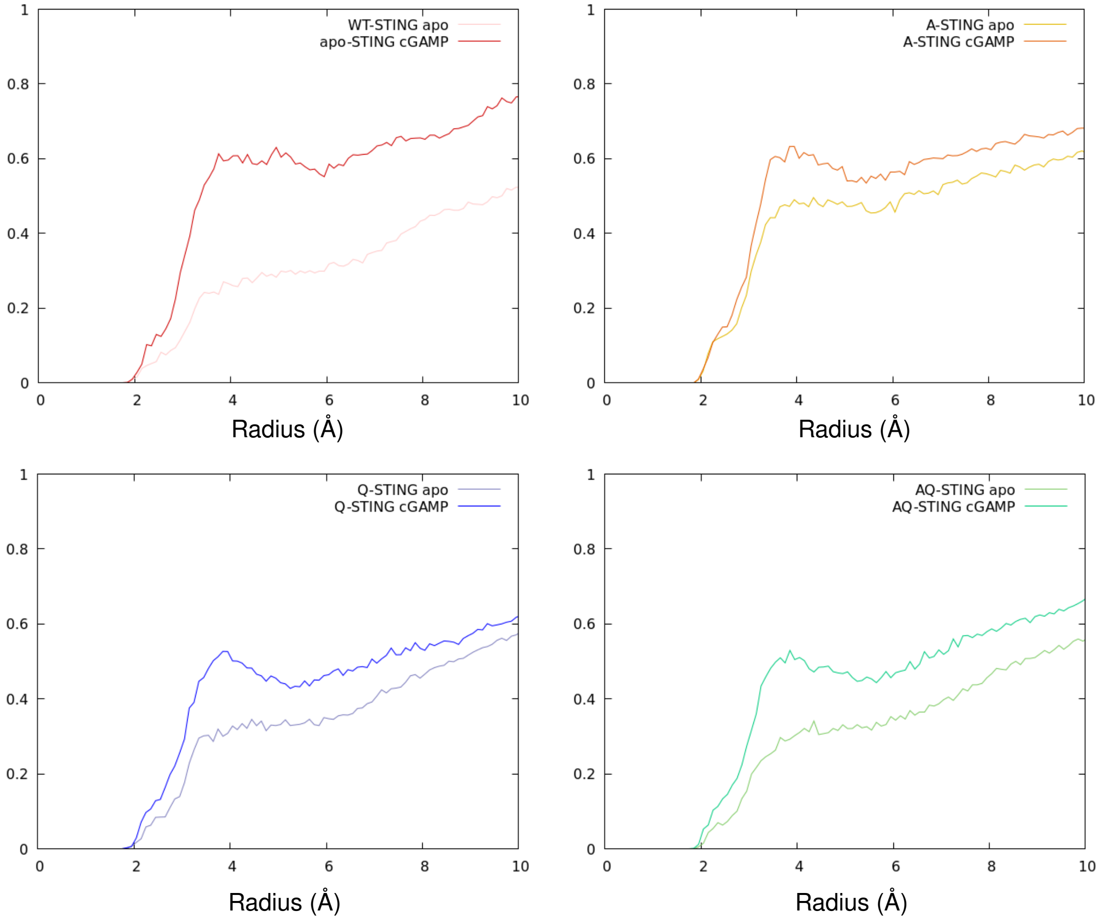


Figure S4 – Probability of presence of water molecules with respect to the radius to the cysteines for each variant.

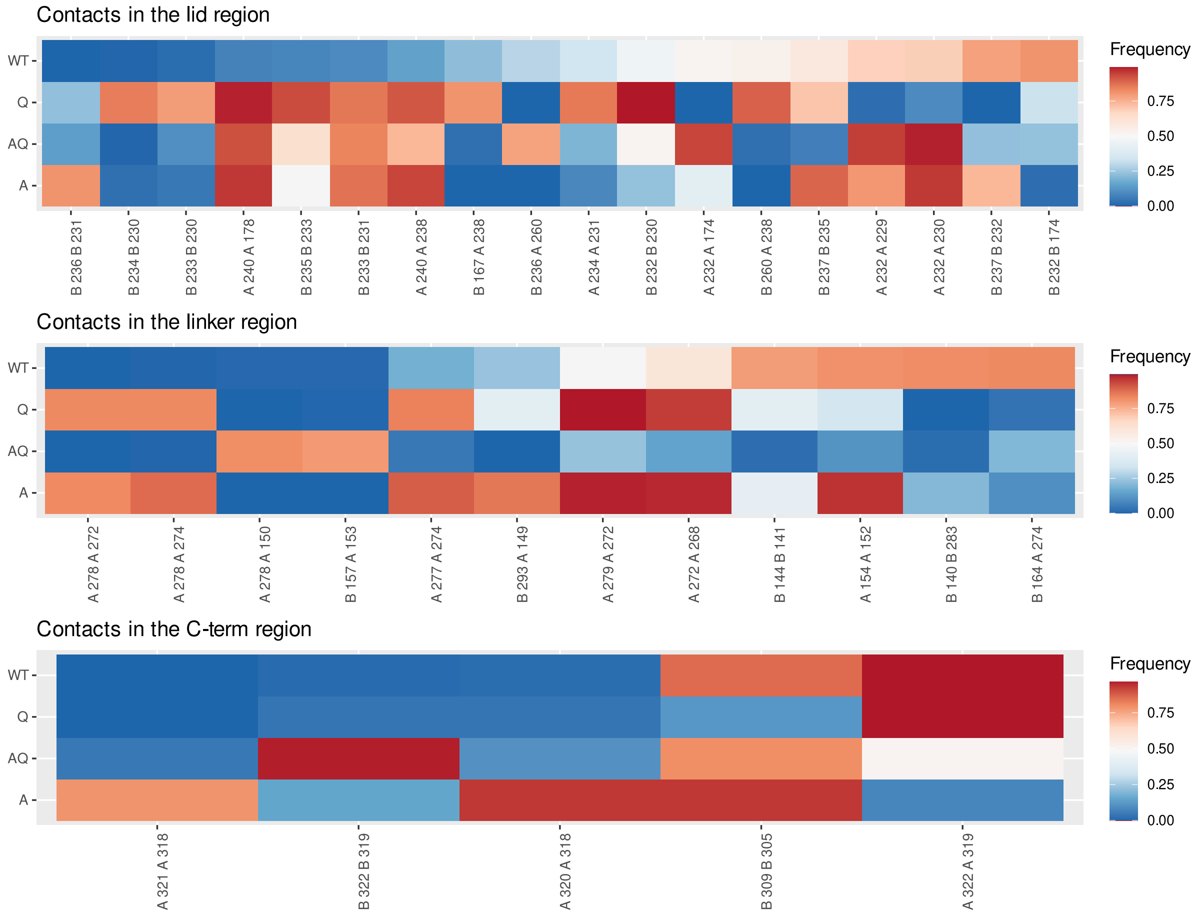


Figure S5 – Differences of contacts frequency among the apo variants, filtered with a threshold of 75% difference with respect to the WT.

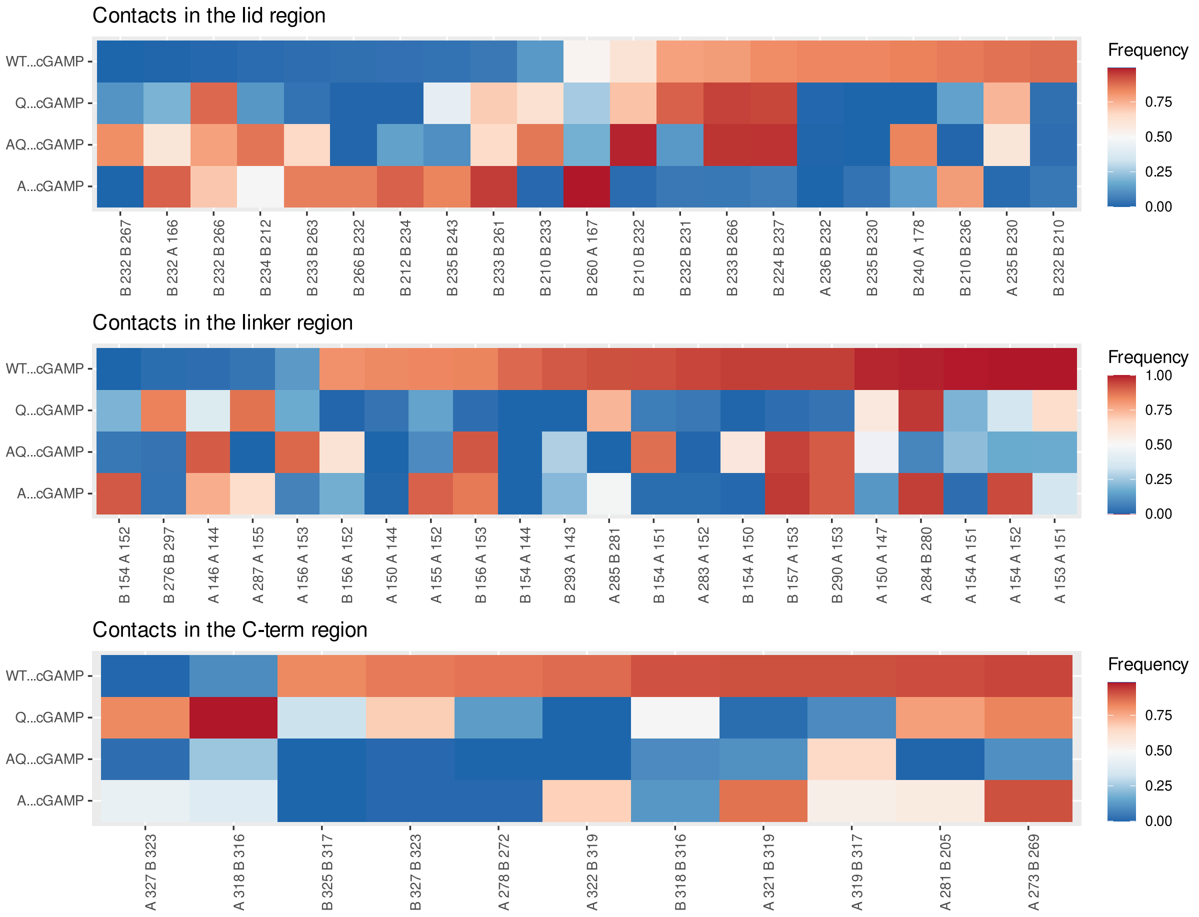


Figure S6 – Differences of contacts frequency among the cGAMP-bound variants, filtered with a threshold of 80% difference with respect to the WT.

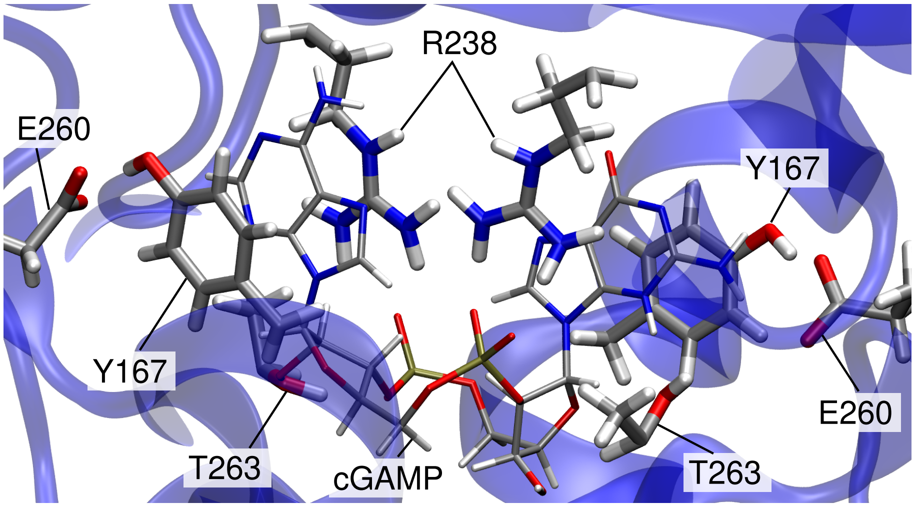


Figure S7 – Organization of the binding site of the Q variant in interaction with cGAMP.

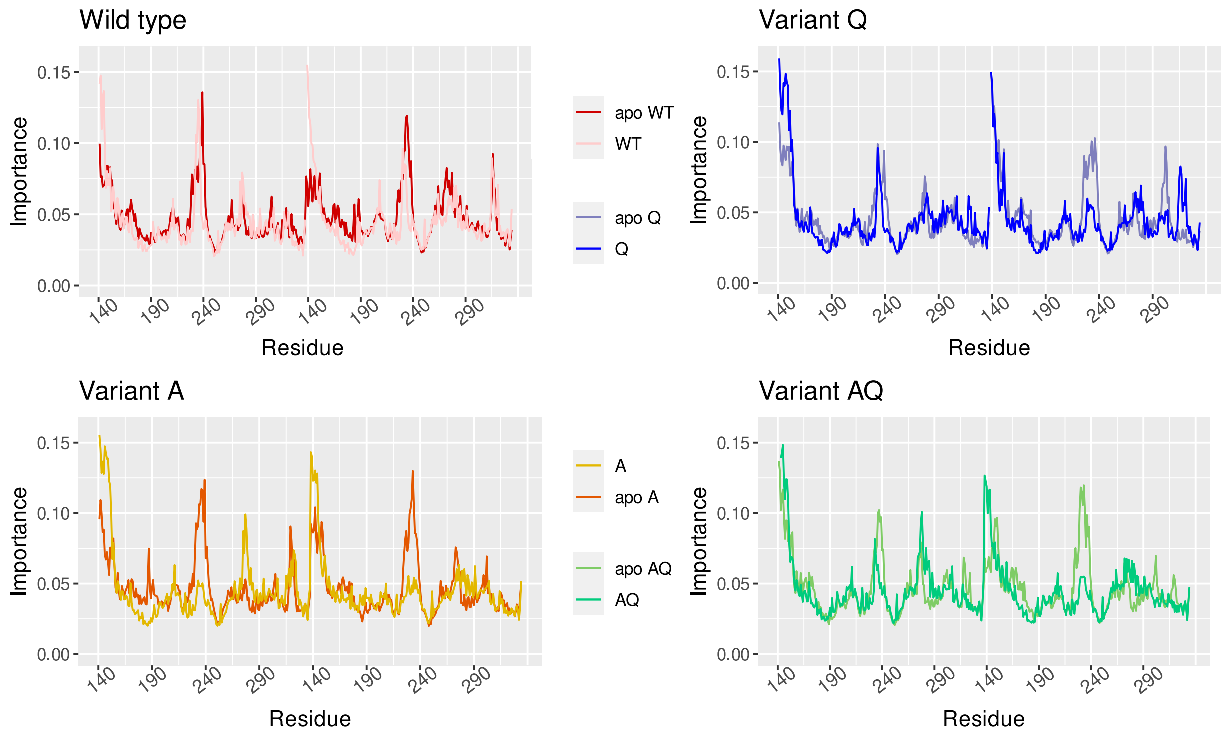


Figure S8 – Flexibility profiles for the apo and bound states of each variant separately.
